# Installation and imaging of thousands of minirhizotrons to phenotype root systems of field-grown plants

**DOI:** 10.1101/2021.09.13.460133

**Authors:** Ashish B. Rajurkar, Scott M. McCoy, Jeremy Ruhter, Jessica Mulcrone, Luke Freyfogle, Andrew D. B. Leakey

## Abstract

**Background:** Roots are vital to plant performance because they acquire resources from the soil and provide anchorage. However, it remains difficult to assess root system size and distribution because roots are inaccessible in the soil. Existing methods to phenotype entire root systems range from slow, often destructive, methods applied to relatively small numbers of plants in the field to rapid methods that can be applied to large numbers of plants in controlled environment conditions. Much has been learned recently by extensive sampling of the root crown portion of field-grown plants. But, information on large-scale genetic and environmental variation in the size and distribution of root systems in the field remains a key knowledge gap. Minirhizotrons are the only established, non-destructive technology that can address this need in a standard field trial. Prior experiments have used only modest numbers of minirhizotrons, which has limited testing to small numbers of genotypes or environmental conditions. This study addressed the need for methods to install and collect images from thousands of minirhizotrons and thereby help break the phenotyping bottleneck in the field.

**Results:** Over three growing seasons, methods were developed and refined to install and collect images from up to3038 minirhizotrons per experiment. Modifications were made to four tractors and hydraulic soil corers mounted to them. High quality installation was achieved at an average rate of up to 84.4 minirhizotron tubes per tractor per day. A set of four commercially available minirhizotron camera systems were each transported by wheelbarrow to allow collection of images of mature maize root systems at an average rate of up to 65.3 tubes per day per camera. This resulted in over 300,000 images being collected in as little as 11 days for a single experiment.

**Conclusion:** The scale of minirhizotron installation was increased by two orders of magnitude by simultaneously using four tractor-mounted, hydraulic soil corers with modifications to ensure high quality, rapid operation. Image collection can be achieved at the corresponding scale using commercially available minirhizotron camera systems. Along with recent advances in image analysis, these advances will allow use of minirhizotrons at unprecedented scale to address key knowledge gaps regarding genetic and environmental effects on root system size and distribution in the field.

## Background

Roots play key roles in anchoring plants to their substrate, acquiring resources from the soil, and as a link between the plant and abiotic or biotic components of the belowground environment (Raven and Edwards 2001; Passioura 2002; Dexter 2004; Marschner 2012). As a result, root system size and distribution significantly impacts plant productivity, sustainability and resilience to stress (Lynch 1995) and root traits have been identified as key targets for crop improvement (Lynch, 2007).

Despite the importance of root systems, many key questions related to genetic and environmental variation in their development, productivity and function remain unanswered because being buried in soil makes them very difficult to measure (Watt et al. 2013; Topp et al. 2016; Atkinson et al. 2019; Wasson et al. 2020). There have been important recent advances in root phenotyping that range from rapid phenotyping of many plants under controlled environment conditions to more arduous, often destructive, methods applied to relatively few plants grown under field conditions (Topp et al. 2016, Atkinson et al. 2019,, Tracy et al. 2020, Wasson et al. 2020).

Transparent growth media in containers allows time courses of 2D or 3D root imaging with digital cameras (Miller et al. 2007; Iyer-Pascuzzi et al. 2010, Topp et al. 2013). Greater realism has been achieved while remaining in controlled environment conditions through the use of opaque artificial media (Downie et al. 2012; Nagel et al. 2012; Lobet and Draye 2013) or soil in pots and other imaging modalities, including X-ray computed tomography (X-ray CT) (Mooney et al. 2012), magnetic resonance imaging (MRI) (van Dusschoten et al. 2016, Rogers et al. 2016), positron emission tomography (PET) (Garbout et al. 2012), neutron radiography and charged-coupled device (CCD)/complementary metal-oxide semiconductor (CMOS) based thermoacoustic imaging (Wasson et al. 2020).

In the field, the trade-off between the realism of the growing environment and the rate or scale of data collection has proven hard to break, with studies of significant genetic diversity in the field most often being limited to destructive measurement of the root crown towards the end of the growing season (Trachsel et al. 2011; Das et al. 2015). Deep roots play a pivotal role in plant function, including acquisition of nutrients and water (Liedgens and Richner, 2001).

Assessing the size and depth distribution of the entire root system of field-grown crops is still only possible with three technologies that have been in use for decades. First, soil cores can be collected and roots washed out in different layers through the soil profile, but this is destructive and very laborious (e.g. Bolinder et al. 1997; Lei et al. 2021). The core-break method (Drew and Saker 1980) can be used to screen tens of genotypes (Wasson et al. 2014) and recently has been made more efficient through innovations in image acquisition and analysis (Wasson et al. 2016), but it also requires destructive sampling at the end of the field season. Third, minirhizotrons can be used to non-destructively image the size and depth distribution of root systems at different times in the growing season at tens of locations within an experimental field (e.g. Bates 1937; Brown and Upchurch 1987; Gray et al. 2013, 2016). Minirhizotrons installed at an angle of 30-45° from vertical are most commonly recommended for use in an open field setting because vertical tubes can cause artifacts in root distribution (Bragg et al. 1983; Johnson et al. 2001; Polomski and Kuhn, 2002). Horizontal tubes have valuable applications, but require trenching to allow installation, and tend to be used in specialized facilities (e.g. Wahlstrom et al. 2021). Therefore, angled minirhizotrons were the focus for developing new methods of rapid installation suitable to general use in crop field trials.

This study aims to facilitate a breakthrough in high-throughput phenotyping of root systems in the field by developing and refining methods that allow thousands of minirhizotrons to be deployed in a single field experiment. This is a significant challenge because minirhizotron installation cannot start until after a crop is sown, otherwise the portion of the minirhizotron tube protruding above the soil surface would interfere with the operation of a mechanical planter. In addition, installation of access tubes must be complete before the crop has germinated and grown to a stage where installation would damage plants. In a field setting, this period lasts only a few weeks. The unpredictability of weather patterns in many locations means that field activities are unlikely to be possible every day even if installation is possible under a range of soil moisture contents.

Minirhizotron tubes can be installed after excavating holes with hand augers (Arnaud et al. 2019), hand held power augers (Kloeppel and Gower 1995), and augers or hydraulic soil corers mounted to vehicles, including tractors (Gray et al. 2013). All of these methods of tube installation have been applied for several decades (Brown and Upchurch 1987; Box et al. 1989). Method development has focused heavily on the best material to use for minirhizotron tubes and ensuring good quality installation i.e. minimal soil disturbance and close contact between the soil and outer surface of the tube throughout its length (Johnson et al. 2001). While this is essential, the scale of experiments has remained limited, with only tens of minirhizotrons being installed in each experiment, even when vehicle/mounted corers were used for installation in relatively easy to access agricultural systems (e.g. Box et al. 1989; Cheng et al. 1990; Murphy et al. 1994; Gray et al. 2013; Gray et al. 2016; Xu et al. 2020). This study built on 10 years of experience using traditional approaches to install minirhizotron tubes. Methods for using tractor-mounted hydraulic corers were modified and accelerated. The use of multiple systems in parallel to scale-up operations was also demonstrated. Finally, we tested if currently available minirhizotron cameras can be used to collect images in a reasonable timeframe when the scale of an experiment increases to use thousands of minirhizotrons.

## Results and Discussion

Methods were successfully developed to install and collect images from 1508 - 3038 minirhizotron tubes in a single experiment in each of three successive growing seasons (Tables 1 and 2; https://www.youtube.com/watch?v=kciMoWH-fFk). The commercially available tractor-mounted hydraulic corer that was used is designed for sampling vertical soil cores (Figure 1). It was modified with a telescoping steel laydown bar to fix the angle of the corer at 30° from vertical (Figure 1). This was used to bore a ∼152 cm-long hole in the ground of the same diameter as the cellulose acetate butyrate (CAB) minirhizotron tube. To avoid soil compaction and accelerate installation, multiple short cores were extracted in succession from each hole. The coring tube was removed after each short core was collected and replaced with a new coring tube. This allowed the next core to be excavated while the soil from the previous coring tube was collected into a bucket. This approach was especially important when soils were wet because it greatly reduces how often the soil core becomes stuck inside the coring tube. Wet soil blocking a series of coring tubes can completely halt minirhizotron installation, and must be avoided. For the same reason, slotted coring tubes that were easier to clean out and coring bits with increasingly small internal diameters (i.e. “quick relief” bits) were used when soils were wetter. After boring the hole, the minirhizotron tube was inserted by placing the PVC cap over the tube and pushing down on it with the base of the soil tube adapter of the hydraulic system. All excavated soil was transported to the boundary of the field in order to avoid altering soil properties near the minirhizotron tubes. The PVC plastic cap covered the portion of the tube protruding from the soil completely, thereby serving to block light transmission that could otherwise alter root growth (Levan et al. 1987; Johnson et al. 2001) and blocking rain from entering the tube.

**Table 1:** Year, Tubes installed, days required and efficiency of installation method

| Year | Planting Date | Beginning of Tube Installation | End of Tube Installation | Plots with Tubes | Tubes Installed | Average Tubes per Plot | Installation Days | Average Tractors Used Per Day | Operators per tractor | Total Worker days | Average Tubes Installed per Worker Hour | Average Tubes Installed per 8 hours per Tractor |
| --- | --- | --- | --- | --- | --- | --- | --- | --- | --- | --- | --- | --- |
| 2017 | 5/31/2017 | 6/5/2017 | 6/13/2017 | 754 | 1508 | 2.00 | 7 | 4 | 2 | 56.0 | 3.37 | 53.9 |
| 2018 | 4/29/2018 | 4/30/2018 | 5/29/2018 | 950 | 3015 | 3.17 | 13 | 3.333 | 2 | 86.7 | 4.35 | 69.6 |
| 2019 | 5/18/2019 | 5/20/2019 | 6/7/2019 | 783 | 3038 | 3.88 | 9 | 4 | 2 | 72.0 | 5.27 | 84.4 |

**Table 2:**
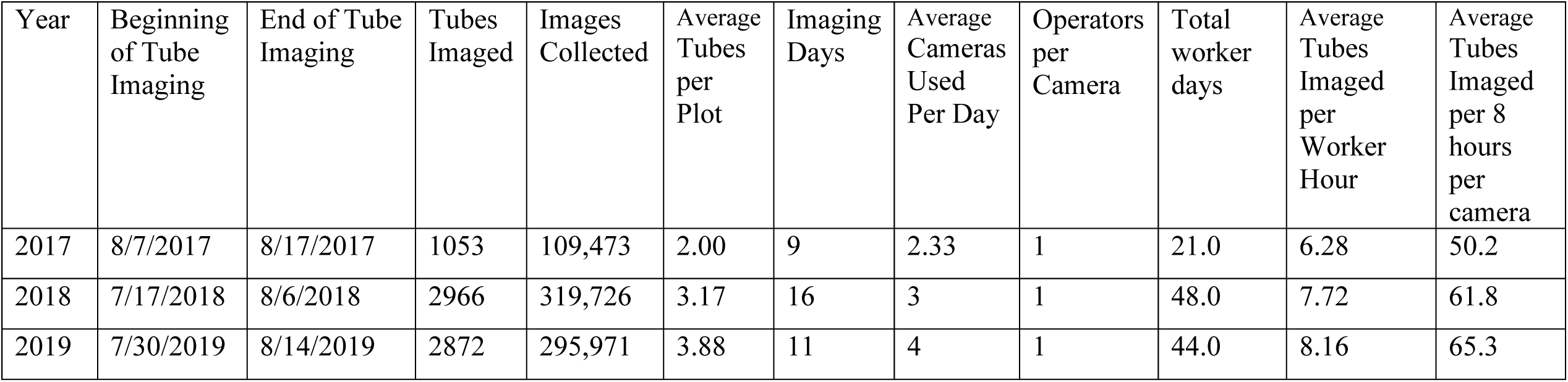
Year, Tubes imaged, images collected, days required and efficiency of imaging

| Year | Beginning of Tube Imaging | End of Tube Imaging | Tubes Imaged | Images Collected | Average Tubes per Plot | Imaging Days | Average Cameras Used Per Day | Operators per Camera | Total worker days | Average Tubes Imaged per Worker Hour | Average Tubes Imaged per 8 hours per camera |
| --- | --- | --- | --- | --- | --- | --- | --- | --- | --- | --- | --- |
| 2017 | 8/7/2017 | 8/17/2017 | 1053 | 109,473 | 2.00 | 9 | 2.33 | 1 | 21.0 | 6.28 | 50.2 |
| 2018 | 7/17/2018 | 8/6/2018 | 2966 | 319,726 | 3.17 | 16 | 3 | 1 | 48.0 | 7.72 | 61.8 |
| 2019 | 7/30/2019 | 8/14/2019 | 2872 | 295,971 | 3.88 | 11 | 4 | 1 | 44.0 | 8.16 | 65.3 |

**Figure 1:**
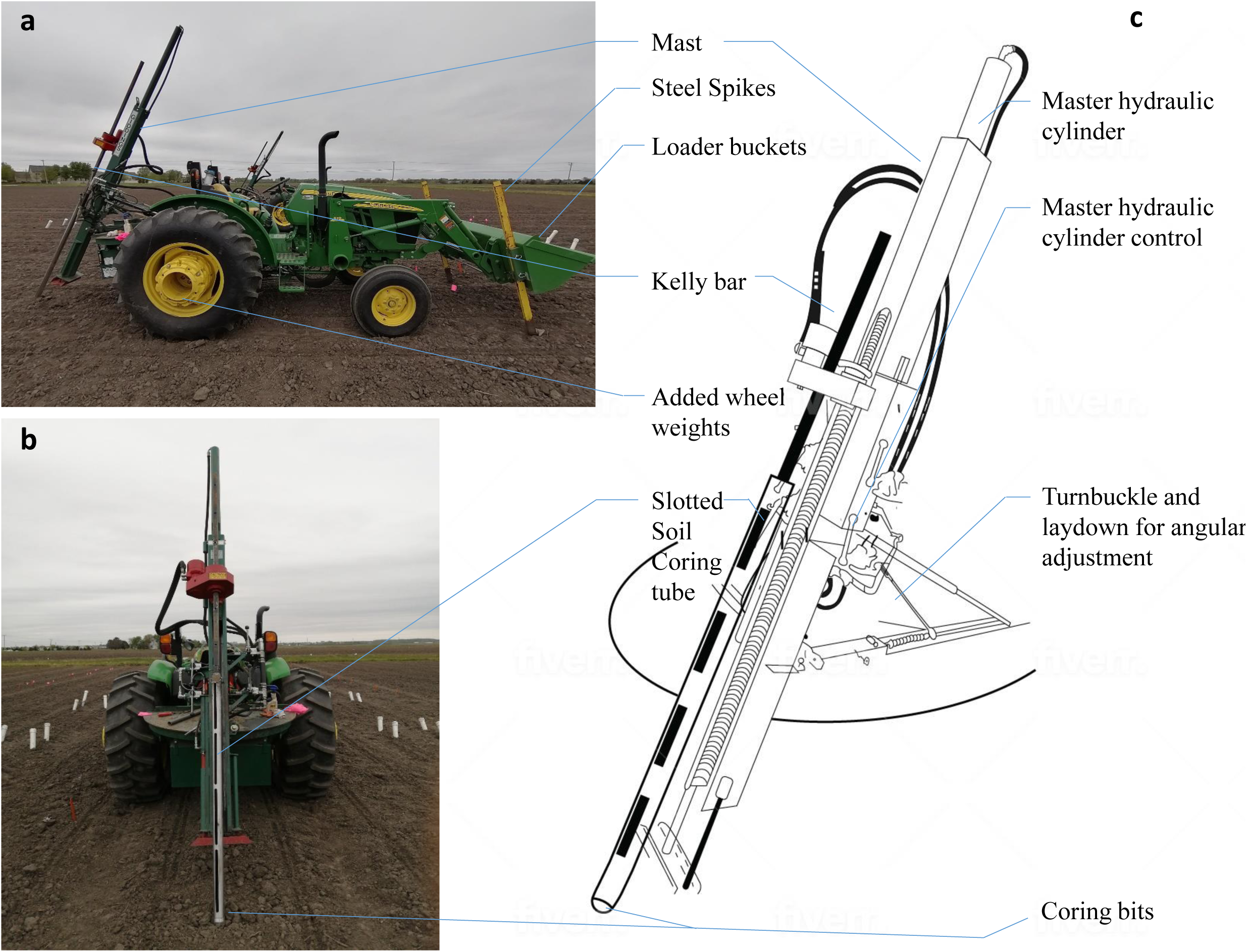
Tractor-mounted Giddings probes shown as: a) side view photograph; b) rear view photograph; and c) diagram of key components and modifications.

The forces required to bore out the hole and insert the minirhizotron tube are considerable, and can be sufficient to lift the tractor off the ground and also push it forwards. If this happens, it can cause the minirhizotron tube to bend and not have good contact with the soil down its entire length. It can also cause damage to equipment and can slow down installation. Therefore, 350 kg of weights were added to the rear wheels of the tractor (Figure 1). In addition, custom-made steel spikes were attached to the front loader so that they could be inserted up to 50 cm into the soil as a physical brake against forward motion during installation.

The quality of root observations made in minirhizotrons is strongly influenced by the method of installation (Cai et al. 2016). The main goal is to make good contact between the surrounding soil and the entire length of the tubes. This requires straight holes to be bored with precision (Johnson et al. 2001). This was aided by the use of a tractor-mounted hydraulic corer with steel coring tubes and tractor weights/stabilization. If this was not achieved, excessive root proliferation would be observed in gaps between the soil and tube surface, as has been reported in some prior occasions (Volkmar 1993; Joslin et al. 1999; Rewald and Ephrath, 2013). However, this was not the case and roots were distributed along the tube in accordance with expectations (Figure 2; Jackson et al. 1996; Black et al. 2017). In addition, radial soil cracks can form in surface soil around the tube causing unwanted changes in root distribution (Rewald and Ephrath, 2013), but this was not observed in the current study.

**Figure 2:**
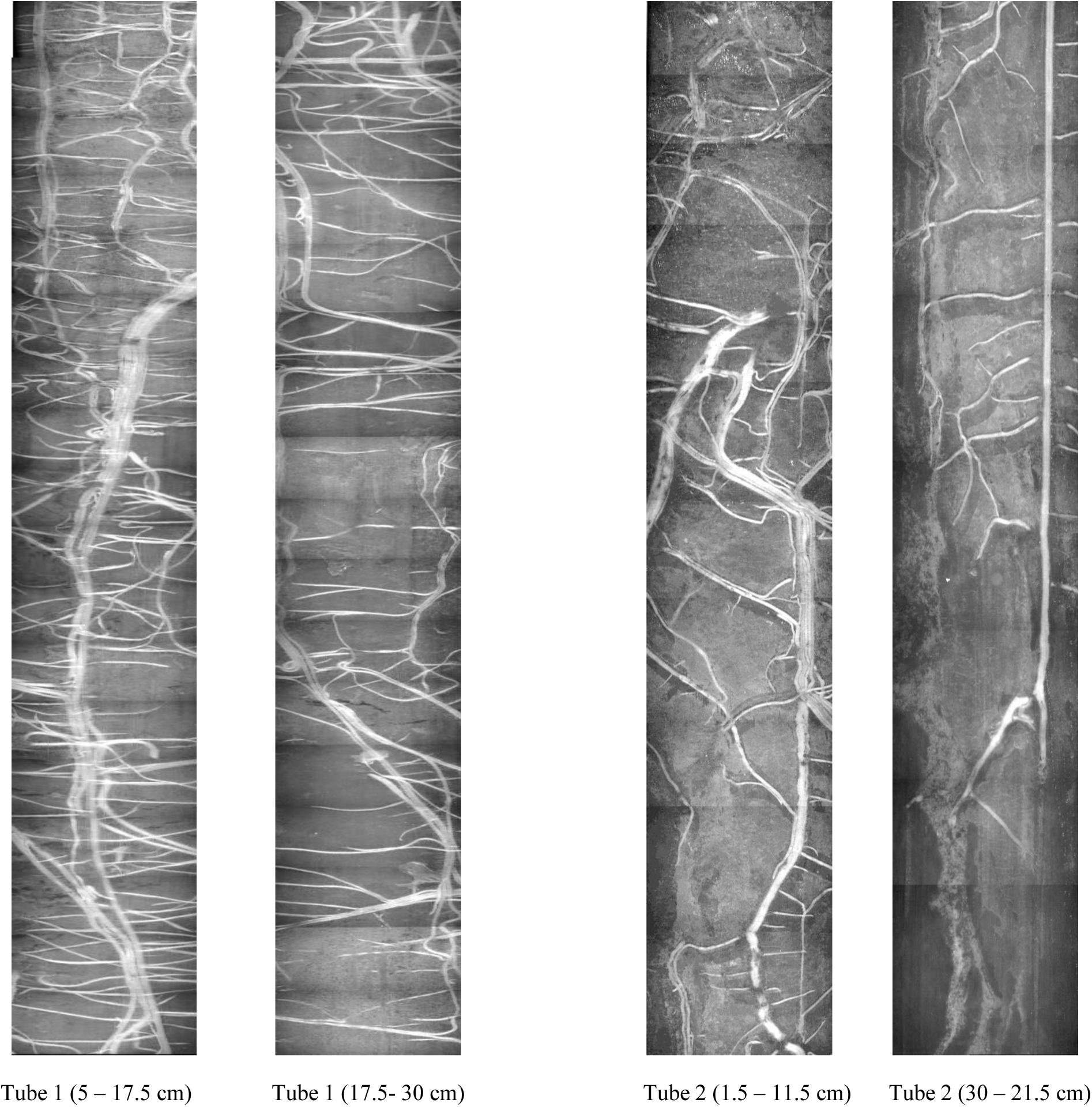
Examples of images from portions of the soil profile from two example minirhizotron tubes showing variation in the root system size and distribution at shallow to moderate depths. Individual images with dimensions of 18 × 13 mm were stitched together using Adobe Photoshop.

Progressive refinement of methods to produce the technique described above allowed the average rate of tube installation per worker per hour to increase from 3.37 in 2017 to 4.35 in 2018 and 5.27 in 2019 (Table 1). This translated into each tractor installing 53.9 to 84.4 tubes on average per day in a given year (Table 1), although a single tractor regularly succeeded in installing >100 tubes in a day. On a typical day, four tractors with hydraulic corers were operated in parallel by a team of eight people i.e. two people per tractor. In the final year, this allowed 3038 minirhizotron tubes to be installed in a single experiment (Table 1; Figure 3) over 9 days of work that were completed within 17 days after sowing. The length and angle of tube protruding from the soil was quickly and easily measured after installation. The tubes were deployed in 783 plots, with 3.88 tubes per plot on average, allowing root system size and distribution to be assessed across hundreds of genotypes of maize.

**Figure 3:**
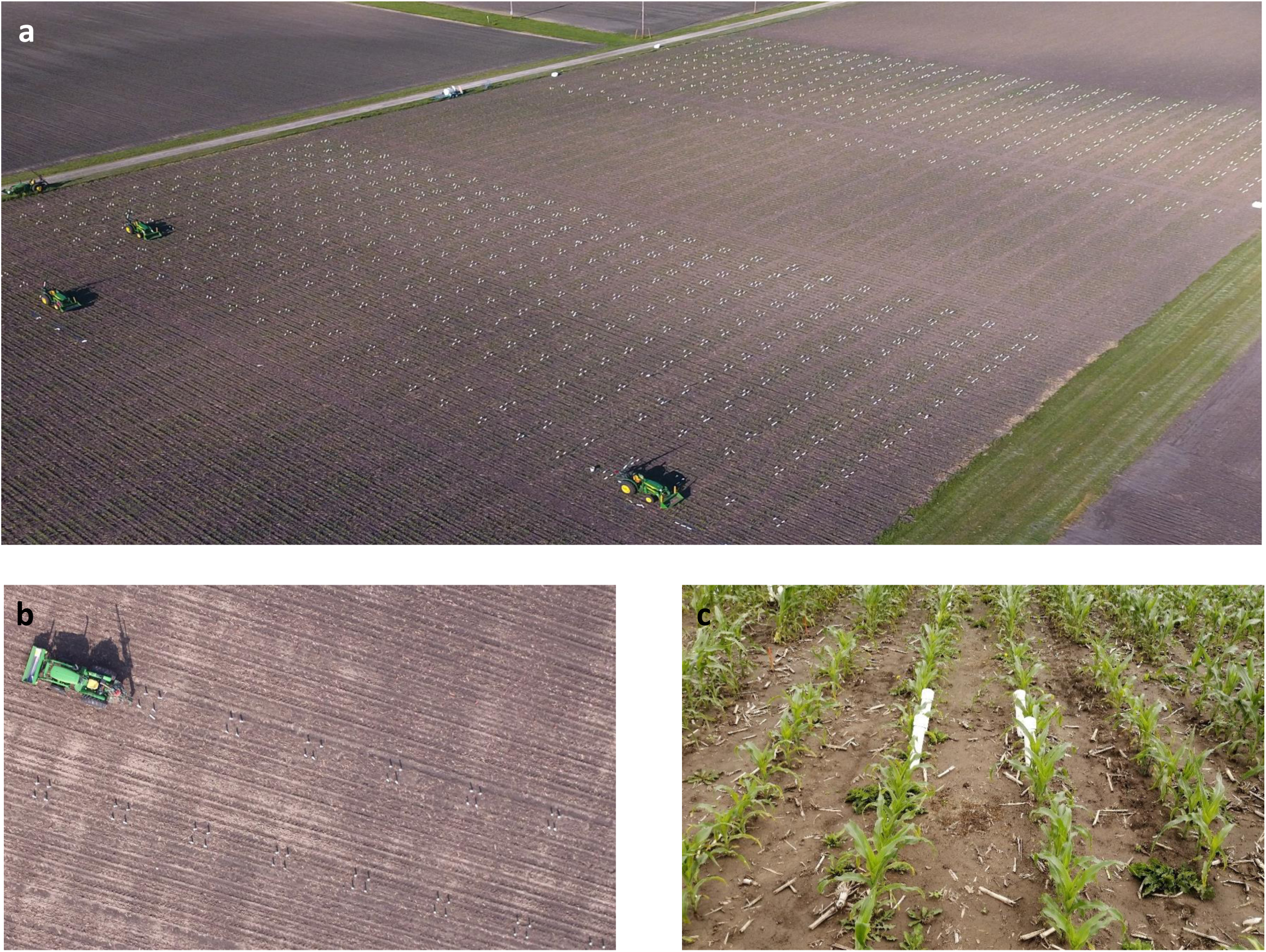
Large scale minirhizotron installation in 2019, demonstrating capacity of installation with the presented method, as a) aerial photograph of full experiment b) aerial photograph of portion of the experiment showing four minirhizotron tubes per subplot c) close up from ground of a single subplot showing PVC caps on tubes and normal plant growth at a later growth stage.

Operating at such a large scale (i.e. >5.5 km of total minirhizotron tubing in a single experiment) requires logistical planning to support the storage, transport and distribution of materials. Minirhizotron tubes, minirhizotron caps and drilling supplies were transported to the edge of the field by forklift and then the materials needed for a day’s work were stored in the front-loader bucket of the tractor, saving considerable manual labor.

Increasing the scale of minirhizotron installation to such a large degree potentially means other aspects of using this root phenotyping approach become unfeasible. Fortunately, tools for automated analysis of images from minirhizotrons are becoming available (Zeng et al. 2008; Svane et al. 2019; Xu et al. 2020). But, image acquisition must also be feasible in a reasonable timeframe. We demonstrated that with four commercially available minirhizotron cameras being used in parallel, the large number of minirhizotrons that were installed could be imaged within 9-16 days when starting immediately after anthesis. This represents a period in time when prior experiments indicated that maize root system size and depth has reached its maximum and is stable prior to the onset of senescence at the end of the growing season (Figure 4). The average number of tubes imaged per day by each camera operator increased from 50.2 in 2017, to 61.8 in 2018, and 65.3 in 2019 (Table 2). In 2017, only 70% of the installed tubes were imaged because drought and heat during and immediately after planting resulted into poor establishment of experimental plots and the plots with poor establishment were excluded from imaging.

**Figure 4:**
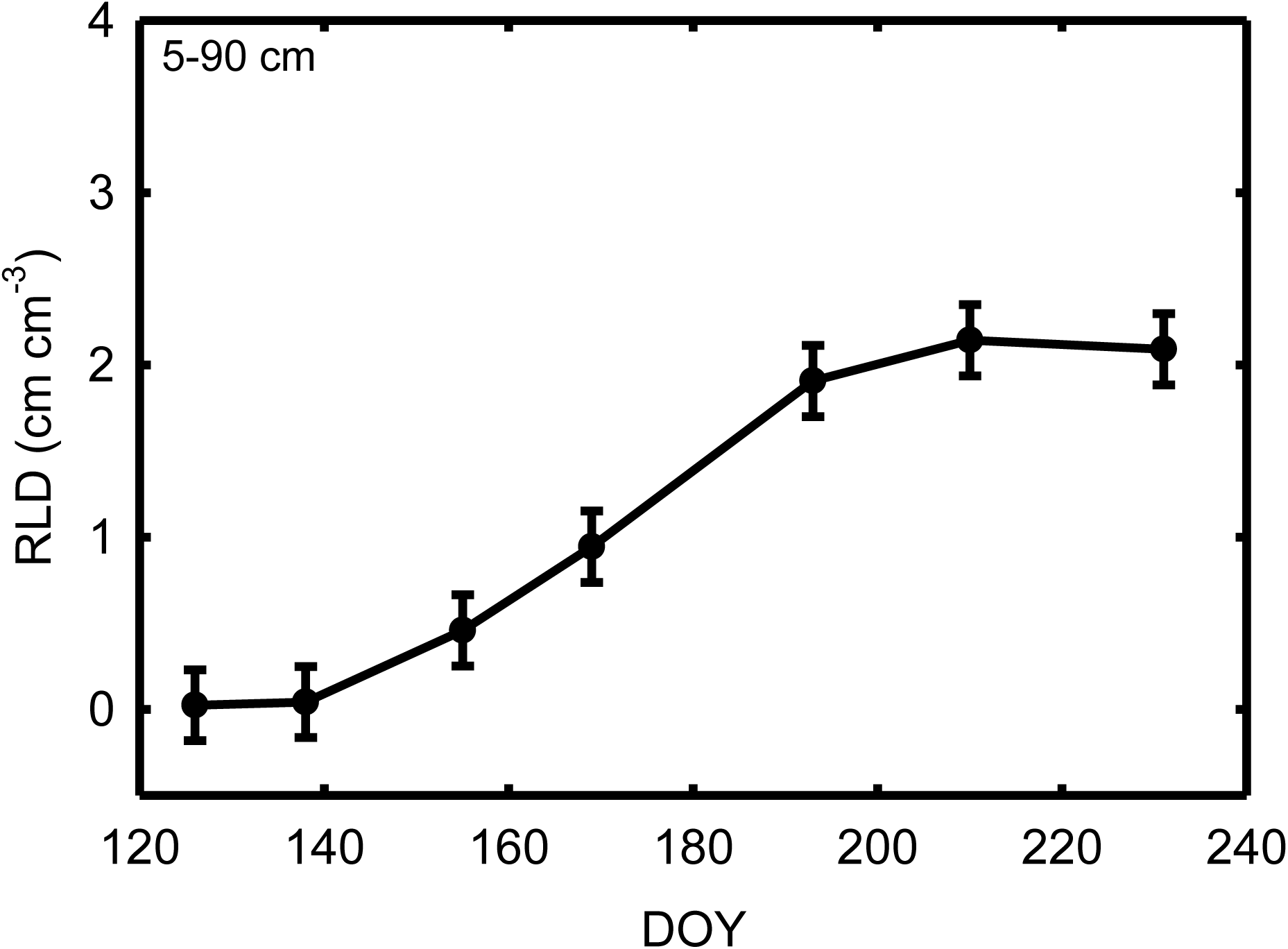
Total root length density (RLD; cm cm-3) of maize at depths of 5-90 cm on mutliple days of year (DOY). Data was collected using the methods of Gray et al. (2016), including manual analysis of all images. Data shown are lsmeans ± standard errors.

This approach means that phenotyping genetic and/or environmental variation in final root system size and distribution is immediately possible. While that does address an important knowledge gap, it would still not be feasible to collect data across all minirhizotrons at other periods of the growing season when the root system is growing at a rapid rate. That limits the ability to test for variation across time in the effect of environmental treatments or genetic variation (Gray et al. 2013, 2016). And, it prevents rates of root system growth (e.g. Figure 4) from being measured, which is significant because data on genetic variation in early-season rates of above-ground growth have proven very valuable in predicting end-of-season biomass production (Varela et al. 2021). Accelerating the rate of image acquisition could be achieved by: (1) a robot or robots that move around the field visiting minirhizotrons and collecting data; (2) cameras that can acquire images automatically when installed permanently in each minirhizotron, or after being moved from minirhizotron to minirhizotron (Childs et al. 2020; Rahman et al. 2020; Thomsen and Jensen 2020); (3) or by a large team of people using many cameras in parallel. All of these options will require further innovation and/or commercialization to make the necessary equipment available and affordable.

## Conclusion

The methods demonstrated in this study represent a substantial increase in the scale of minirhizotron experimentation, and open up the possibility of using the method in studies that use quantitative genetic techniques to discover the phenotype-to-genotype associations as well as trait relationships that are current major knowledge gaps in root biology (Watt et al. 2013; Topp et al. 2016; Atkinson et al. 2019; Wasson et al. 2020).

## Methods

### Field site

Diverse populations of 369 inbred maize lines (2017) and 700 hybrid maize lines (2018 and 2019) were planted in replicated blocks on the experimental farms of the University of Illinois at Urbana-Champaign with a precision planter. Each subplot containing a single genotype had four rows with length of 3.65 m, row spacing of 0.76 m, and plant spacing within a row of 0.18 m.

### Minirhizotron materials

Cellulose acetate butyrate (CAB) minirhizotron tubes of 183-cm (6 ft.) length, 5.715-cm diameter, wall thickness of 0.3175 cm, and internal diameter of 5.3975 cm were used (Exelon by Thermo, Georgetown, Delaware, USA). Tubes were sealed at the bottom with low-density polyethylene cap plugs (product # CCF-2 1/4-10-14, Caplugs, Buffalo, NY, USA; 5.7150 cm outer diameter) and glued with adhesive (Loctite Instant mix) and all-purpose non-toxic sealant (GE All Purpose Silicone 1) to avoid water infiltration from the soil into the tube. A small indexing hole was drilled near the top of the tube using a guide to maintain consistent position (Bartz, CA, USA) which helps in locking the camera to the tube during imaging. Tubes were stored in batches of 300 in collapsible containers (Uline product # H-1732), which had wooden separators for easy stacking and to avoid shifting during transport to the field by forklift.

Minirhizotron caps that prevented rain from entering the tube and excluded light from being transmitted below-ground by the tube were constructed from 6.35-cm diameter, schedule 40 PVC pipe. Each cap was 31 cm long, with one end closed by a socket cap (product # 447-25, Lasco Fittings Inc. Brownsville, TN, USA) and the other end cut at an angle to rest upon the soil surface. The caps were stored and transported in polypropylene bags with forklift straps that are designed for transport of building materials.

### Installation equipment

Four sets of hydraulic soil coring probes (Model 15GSRTS, Giddings Machine Co., Windsor, CO, USA) mounted to small tractors (Model 5065E with 512 front loader, John Deere, Moline, IL, USA; Figure 1) were used to bore holes, insert minirhizotron tubes, and extract minirhizotron tubes. The tractors were fitted with 350 kg rear-wheel weights. Steel bars 2.5 cm thick, 10 cm wide and 183 cm long were custom fabricated and mounted to the front loader bucket to act as a physical brake against the forward force produced by the corer during installation (Figure 1a). The spikes were tapered to aid insertion into the soil as a physical break and had several sets of mounting holes to allow length adjustment during transport and installation.

Each hydraulic corer was modified to include a telescoping tubular steel bar that maintained the mast at the desired angle of 30° from vertical during installation but still allowed the mast to be “laid down” when tubes are not being installed. A heavy-duty Kelly bar (KB-308) with a 5.715 cm soil tube adapter was used to mount the coring tube. A steel soil coring tube of long slotted or solid wall type (152 cm (5 ft.) length and 5.715 cm outer diameter) was fit with sharpened coring bits. This tube exactly matched the outer diameter of the minirhizotron tubes. The internal diameter of the bit was progressively reduced as tubes needed to be installed in wetter and heavier soils: from standard (4.7625 cm) in dry soils, to quick-relief (4.6037 cm), heavy quick relief (4.445 cm) and extra heavy quick relief (4.2735 cm) in wet, heavy clays. The use of slotted type soil corer tubes helped in avoiding compaction. Also, if coring tubes became clogged with soil, pry bars were inserted into the slit to clear the blockage.

### Installation technique

Two to four minirhizotron tubes were installed in each genotype subplot. The minirhizotrons were located centrally in each 4-row plot (Figure 3). The locations were predetermined and marked with flags. The tractor was driven into the field straddling the two rows of crop where minirhizotron tubes were to be installed and stopped over the initial installation position. The front loader was lowered to insert the steel bars into the soil as a physical brake. The mast of the hydraulic corer was rotated on the turntable such that the coring tube would enter the soil directly within the row of crops. To avoid compression of soil, repeated short cores were taken with the use of multiple soil corer tubes to make each ∼152-cm long hole (https://www.youtube.com/watch?v=kciMoWH-fFk). The soil from each excavated core was collected in a waste bucket and dumped at the edge of the field to avoid mixing of subsoil with topsoil in experimental plots. After the hole was cored out, a cap was placed over the minirhizotron tube and it was pushed into the hole by placing applying pressure from the hydraulic system to the top of the cap with the soil tube adapter. It was important to maintain steady, low pressure from the hydraulic system to ensure a smooth installation of tubes and to avoid bending or breakage. After the tube was installed, the mast of the hydraulic corer was swung over to allow installation of the next tube in the other middle row of crops, or driven forward to the next position for tube installation in the same row. After all minirhizotron tubes in a given plot were installed, the tractor was driven to the next plot and installation continued. The tractors always drove in a south to north direction so that all tubes had the same orientation. If all the tubes in a given range of plots were not completed within a day, the tractor was left in the field and installation continued from that point the next day. This minimized traffic through the plots to a single pass of the tractor in all cases.

### Minirhizotron angle and depth

A digital clinometer app on cell phones was used to measure the angle of installed minirhizotron tubes and the length of the tube protruding from the soil was measured with a measuring tape in order to subsequently calculate the vertical depth distribution of roots. Tubes were typically inserted to a depth of 152 cm, but on rare occasions (< 0.1% of all tubes) a tube could not be fully inserted. If tubes were installed to ≥75 % of full length they were trimmed to the appropriate height above-ground and images collected from the portion of the tube that could be used. If tubes were installed < 75 % of full length they were trimmed to the soil surface and dropped from the experiment.

### Image Collection

Four minirhizotron video cameras (BTC100x, Bartz Technology Corp, Carpinteria, CA, USA) with incandescent illumination were used in parallel to capture digital color images of roots. The camera was attached to an indexing handle and controlled with a laptop computer-based image capture system (I-CAP, Bartz Technology Corp, Carpentaria, CA, USA). Prior to imaging roots, a reference grid (1 × 1 mm grid) was imaged after adjusting zoom, light intensity and the image perimeter area (2 cm wide) to standard settings that were maintained throughout data collection. The laptop and its 12V battery power source were kept in a wheelbarrow, which made it easier for a single operator to move from plot to plot within the field over long periods of image collection.

Prior to imaging, each minirhizotron tube was checked for water and was cleaned with a long-handled swab for dust and water condensation inside the inner surface of the tube. Images were collected from the upward facing surface of the tube at increments of 1.3 cm down the length of the tube, and at a speed of 1200 milliseconds per index handle location. The quality of images was monitored on the laptop. When images were identified as blurry or out of focus, that minirhizotron was re-imaged. The resulting images had a resolution of 28 µm per pixel and were each 640 × 480 pixels (18 × 13.5 mm) in size. Collectively, they record the distribution of root length across a range of depths of the soil profile (e.g. Figure 2).

### Minirhizotron tube removal

At the end of the growing season, after all sampling was complete, the same tractor-mounted Giddings were used to pull the minirhizotron tubes out of the ground by attaching a Sentek access tube extraction tool (Fondriest Environmental – Fairborn, Ohio, USA) and chains bolted to the Kelly bar. This procedure was not particularly time sensitive and was typically completed at a rate of ∼ 300 tubes per day with two operators for each of two tractors.

## Funding

The research was funded by the Plant Genome Research Program at National Science Foundation under award IOS-1638507 and by the Advanced Research Projects Agency-Energy (ARPA-E), U.S. Department of Energy, under Award Number DE-DE-AR0000661.

## Acknowledgments

We thank Brady Adams, Dylan Allen, Darshi Banan, Jim Berry, Tyler Blackwell, Aya Bridgeland, Setu Chakrabarty, Benjamin Daniels, Brooks Hauser, Benjamin Ihssen, Avinash Karn, Shahbaz Khan, Zac Lumley, Grace McDonough, Jesse McGrath, Adrian Madera, Andrew Mareczko, Mike Masters, Dhruv Mishra, Chris Montes, Askar Mukhanov, Jess Mulcrone, Belen Muniz, Madison Murray, Ndidiamaka Ojiako, Savanna Pflugmacher, Melina Pakey-Rodriguez, Cody Randolph, Peter Schmuker, Lincoln Taylor, Sean Tobin, Ben Thompson, Austin von Perbandt, and Tim Wertin for assistance with field work. We thank Stephen Moose and Sherry Flint-Garcia for supplying seed. We thank Scott Baker and Jared Bear at the Life Sciences Machine Shop for technical assistance. We thank Ivan Baxter and Chris Topp for stimulating discussion on experimental design and assistance with project planning.

## Authors’ Contributions

ADBL conceived the study. SM, JR, JM and LF developed methods and equipment. ABR, SM, JR, JM, LF and ABDL performed experiments. ABR and ADBL wrote the manuscript. All authors read and approved final manuscript.

## Ethics approval and consent to participate

Not applicable

## Consent for publication

Not applicable

## Availability of data and material

Not applicable

## Competing interests

The authors declare that they have no competing interests

## Notes

### Competing Interest Statement

The authors have declared no competing interest.

## References

Arnaud M, Baird AJ, Morris PJ, Harris A, Huck JJ. EnRoot: a narrow-diameter, inexpensive and partially 3D-printable minirhizotron for imaging fine root production. Plant Methods. 2019;15(1):1–9.

Atkinson JA, Pound MP, Bennett MJ, Wells DM. Uncovering the hidden half of plants using new advances in root phenotyping. Current opinion in biotechnology. 2019;55:1–8.

Bates GH. A device for the observation of root growth in the soil. Nature. 1937;139(3527):966–7.

Black CK, Masters MD, LeBauer DS, Anderson-Teixeira KJ, DeLucia EH. Root volume distribution of maturing perennial grasses revealed by correcting for minirhizotron surface effects. Plant and Soil. 2017;419(1):391–404.

Bolinder MA, Angers DA, Dubuc JP. Estimating shoot to root ratios and annual carbon inputs in soils for cereal crops. Agricukture Ecosystems & Environment. 1997;63(1):61–66.

Box Jr JE, Smucker AJ, Ritchie JT. Minirhizotron installation techniques for investigating root responses to drought and oxygen stresses. Soil Science Society of America Journal. 1989;53(1):115–8.

Bragg PL, Govi G, Cannell RQ. A comparison of methods, including angled and vertical minirhizotrons, for studying root growth and distribution in a spring oat crop. Plant and Soil. 1983;73(3):435–40.

Brown DA, Upchurch DR. Minirhizotrons: a summary of methods and instruments in current use. In: Minirhizotron observation tubes - methods and applications for measuring rhizosphere dynamics, edited by H.M. Taylor. Proceedings of a Symposium, December 3, 1986, New Orleans, Louisiana. ASA-special-publication-American-Society-of-Agronomy (USA). 1987;50:15–30.

Cai G, Vanderborght J, Klotzsche A, van der Kruk J, Neumann J, Hermes N, Vereecken H. Construction of minirhizotron facilities for investigating root zone processes. Vadose Zone Journal. 2016;15(9).

Cheng W, Coleman DC, Box Jr JE. Root dynamics, production and distribution in agroecosystems on the Georgia Piedmont using minirhizotrons. Journal of Applied Ecology. 1990:592–604.

Childs J, Defrenne CE, Brice DJ, Woodward J, Holbrook KN, Nettles WR, Taggart M, Iversen CM. SPRUCE high-resolution minirhizotrons in an experimentally-warmed peatland provide an unprecedented glimpse at fine roots and their fungal partners: Supporting data. Oak Ridge National Lab.(ORNL), Oak Ridge, TN (United States); 2020.

Das A, Schneider H, Burridge J, Ascanio AK, Wojciechowski T, Topp CN, Lynch JP, Weitz JS, Bucksch A. Digital imaging of root traits (DIRT): a high-throughput computing and collaboration platform for field-based root phenomics. Plant Methods. 2015;11(1):1–2.

Dexter AR. Soil physical quality: Part I. Theory, effects of soil texture, density, and organic matter, and effects on root growth. Geoderma. 2004;120(3-4):201–14.

Downie H, Holden N, Otten W, Spiers AJ, Valentine TA, Dupuy LX. Transparent soil for imaging the rhizosphere. PLoS One. 2012;7(9):e44276.

Drew MC, Saker LR. Assessment of a rapid method, using soil cores, for estimating the amount and distribution of crop roots in the field. Plant and Soil. 1980;55(2):297–305.

Flint-Garcia SA, Thuillet AC, Yu J, Pressoir G, Romero SM, Mitchell SE, Doebley J, Kresovich S, Goodman MM, Buckler ES. Maize association population: a high-resolution platform for quantitative trait locus dissection. The Plant Journal. 2005;44(6):1054–64.

Garbout A, Munkholm LJ, Hansen SB, Petersen BM, Munk OL, Pajor R. The use of PET/CT scanning technique for 3D visualization and quantification of real-time soil/plant interactions. Plant and Soil. 2012;352(1-2):113–27.

Gray SB, Strellner RS, Puthuval KK, Ng C, Shulman RE, Siebers MH, Rogers A, Leakey ADB. Minirhizotron imaging reveals that nodulation of field-grown soybean is enhanced by free-air CO2 enrichment only when combined with drought stress. Functional Plant Biology. 2013;40(2):137–47.

Gray SB, Dermody O, Klein SP, Locke AM, Mcgrath JM, Paul RE, Rosenthal DM, Ruiz-Vera UM, Siebers MH, Strellner R, Ainsworth EA, Bernacchi CJ, Long SP, Ort DR, Leakey ADB. Intensifying drought eliminates the expected benefits of elevated carbon dioxide for soybean. Nature Plants. 2016;2(9):1–8.

Iyer-Pascuzzi AS, Symonova O, Mileyko Y, Hao Y, Belcher H, Harer J, Weitz JS, Benfey PN. Imaging and analysis platform for automatic phenotyping and trait ranking of plant root systems. Plant Physiology. 2010;152(3):1148–57.

Jackson RB, Canadell J, Ehleringer JR, Mooney HA, Sala OE, Schulze ED. A global analysis of root distributions for terrestrial biomes. Oecologia. 1996;108(3):389–411.

Johnson MG, Tingey DT, Phillips DL, Storm MJ. Advancing fine root research with minirhizotrons. Environmental and Experimental Botany. 2001;45(3):263–89.

Joslin JD, Wolfe MH. Disturbances during minirhizotron installation can affect root observation data. Soil Science Society of America Journal. 1999;63(1):218–21.

Kloeppel BD, Gower ST. Construction and installation of acrylic minirhizotron tubes in forest ecosystems. Soil Science Society of America Journal. 1995;59(1):241–3.

Lei C, Abraha M, Chen J, Su YJ. Long-term variability of root production in bioenergy crops from ingrowth core measurements. Journal of Plant Ecology. 2021;14(5):757–70.

Levan MA, Ycas JW, Hummel JW. Light leak effects on near-surface soybean rooting observed with minirhizotrons. In: Minirhizotron observation tubes - methods and applications for measuring rhizosphere dynamics, edited by H.M. Taylor. Proceedings of a Symposium, December 3, 1986, New Orleans, Louisiana. ASA-special-publication-American-Society-of-Agronomy (USA). 1987;50:89–98.

Liedgens M, Richner W. Minirhizotron observations of the spatial distribution of the maize root system. Agronomy Journal. 2001;93(5):1097–104.

Lobet G, Draye X. Novel scanning procedure enabling the vectorization of entire rhizotron-grown root systems. Plant Methods. 2013;9(1):1.

Lynch J. Root architecture and plant productivity. Plant Physiology. 1995;109(1):7.

Lynch JP. Roots of the second green revolution. Australian Journal of Botany. 2007;55(5):493–512.

McMullen MD, Kresovich S, Villeda HS, Bradbury P, Li H, Sun Q, Flint-Garcia S, Thornsberry J, Acharya C, Bottoms C, Brown P. Genetic properties of the maize nested association mapping population. Science. 2009;325(5941):737–40.

Marschner, H., 2012. Marschner’s mineral nutrition of higher plants. Academic press.

Miller ND, Parks BM, Spalding EP. Computer-vision analysis of seedling responses to light and gravity. Plant Journal. 2007; 52(2):374–381.

Mooney SJ, Pridmore TP, Helliwell J, Bennett MJ. Developing X-ray computed tomography to non-invasively image 3-D root systems architecture in soil. Plant and Soil. 2012;352(1-2):1–22.

Murphy JA, Hendricks MG, Rieke PE, Smucker AJ, Branham BE. Turfgrass root systems evaluated using the minirhizotron and video recording methods. Agronomy Journal. 1994;86(2):247–50.

Nagel KA, Putz A, Gilmer F, Heinz K, Fischbach A, Pfeifer J, Faget M, Blossfeld S, Ernst M, Dimaki C, Kastenholz B. GROWSCREEN-Rhizo is a novel phenotyping robot enabling simultaneous measurements of root and shoot growth for plants grown in soil-filled rhizotrons. Functional Plant Biology. 2012;39(11):891–904.

Passioura JB. Soil conditions and plant growth. Plant, Cell & Environment. 2002;25(2):311–8.

Polomski J, Kuhn N. Root research methods. In Y. Waisel, A. Eshel, & U. Kafkafi (Eds.), Plant Roots: The hidden half. 2002;3:295–321.

Raven JA, Edwards D. Roots: evolutionary origins and biogeochemical significance. Journal of Experimental Botany. 2001;52(suppl_1):381–401.

Rahman G, Sohag H, Chowdhury R, Wahid KA, Dinh A, Arcand M, Vail S. SoilCam: A fully automated minirhizotron using multispectral imaging for root activity monitoring. Sensors. 2020;20(3):787.

Rewald, B., & Ephrath, J. E. Minirhizotron techniques. In: Eshel A, Beeckman T, editors. Plant roots: the hidden half. NY:CRC press; 2013. p. 1–15

Rogers ED, Monaenkova D, Mijar M, Nori A, Goldman DI, Benfey PN. X-ray computed tomography reveals the response of root system architecture to soil texture. Plant Physiology. 2016;171(3):2028–40.

Svane SF, Dam EB, Carstensen JM, Thorup-Kristensen K. A multispectral camera system for automated minirhizotron image analysis. Plant and Soil. 2019;441(1):657–72.

Thomsen S, Jensen K. An affordable, fully-automated minirhizotron system for observing fine-root dynamics. InEGU General Assembly Conference Abstracts 2020 (p. 22448).

Topp CN, Iyer-Pascuzzi AS, Anderson JT, Lee CR, Zurek PR, Symonova O, Zheng Y, Bucksch A, Mileyko Y, Galkovskyi T, Moore BT. 3D phenotyping and quantitative trait locus mapping identify core regions of the rice genome controlling root architecture. Proceedings of the National Academy of. 2013;110(18):E1695–704.

Topp CN, Bray AL, Ellis NA, Liu Z. How can we harness quantitative genetic variation in crop root systems for agricultural improvement?. Journal of Integrative Plant Biology. 2016;58(3):213–25.

Trachsel S, Kaeppler SM, Brown KM, Lynch JP. Shovelomics: high throughput phenotyping of maize (Zea mays L.) root architecture in the field. Plant and Soil. 2011;341(1):75–87.

Tracy SR, Nagel KA, Postma JA, Fassbender H, Wasson A, Watt M. Crop improvement from phenotyping roots: Highlights reveal expanding opportunities. Trends in plant science. 2020;25(1):105–18.

van Dusschoten D, Metzner R, Kochs J, Postma JA, Pflugfelder D, Bühler J, Schurr U, Jahnke S. Quantitative 3D analysis of plant roots growing in soil using magnetic resonance imaging. Plant Physiology. 2016;170(3):1176–88.

Varela S, Pederson T, Bernacchi CJ, Leakey AD. Understanding Growth Dynamics and Yield Prediction of Sorghum Using High Temporal Resolution UAV Imagery Time Series and Machine Learning. Remote Sensing. 2021;13(9):1763.

Volkmar KM. A comparison of minirhizotron techniques for estimating root length density in soils of different bulk densities. Plant and Soil. 1993;157(2):239–45.

Wahlström EM, Kristensen HL, Thomsen IK, Labouriau R, Pulido-Moncada M, Nielsen JA, Munkholm LJ. Subsoil compaction effect on spatio-temporal root growth, reuse of biopores and crop yield of spring barley. European Journal of Agronomy. 2021;123:126225.

Wasson AP, Rebetzke GJ, Kirkegaard JA, Christopher J, Richards RA, Watt M. Soil coring at multiple field environments can directly quantify variation in deep root traits to select wheat genotypes for breeding. Journal of Experimental Botany. 2014;65(21):6231–49.

Wasson A, Bischof L, Zwart A, Watt M. A portable fluorescence spectroscopy imaging system for automated root phenotyping in soil cores in the field. Journal of Experimental Botany. 2016;67(4):1033–43.

Wasson AP, Nagel KA, Tracy S, Watt M. Beyond digging: noninvasive root and rhizosphere phenotyping. Trends in Plant Science. 2020;25(1):119–20.

Watt M, Moosavi S, Cunningham SC, Kirkegaard JA, Rebetzke GJ, Richards RA. A rapid, controlled-environment seedling root screen for wheat correlates well with rooting depths at vegetative, but not reproductive, stages at two field sites. Annals of Botany. 2013;112(2):447–55.

Xu W, Yu G, Zare A, Zurweller B, Rowland DL, Reyes-Cabrera J, Fritschi FB, Matamala R, Juenger TE. Overcoming small minirhizotron datasets using transfer learning. Computers and Electronics in Agriculture. 2020;175:105466.

Zeng G, Birchfield ST, Wells CE. Automatic discrimination of fine roots in minirhizotron images. New Phytologist. 2008;177(2):549–57.

